# Bacterial and Archaeal Communities in Recycling Effluents from a Bauxite Flotation Plant

**DOI:** 10.1101/529958

**Authors:** LIU Xinxing, WU Yong-Hong, Xi Liu, Wu Han-yan, XIE Jianping, Guohua Wang, QIU Guan-zhou, HUO Qiang

## Abstract

Recycling effluent has become a bottleneck and an environmental risk associated with the regular production of bauxite via flotation and the sustainable development of the aluminum industry in China. To find a practical direction for biotreatment, the bacterial and archaeal communities in recycling effluents containing concentrate and tailings from bauxite flotation plants were investigated by a metagenomic sequencing method in association with the evaluated geochemical properties. The results showed that *Paracoccus*, *Desulfomicrobium*, *Exiguobacterium*, *Tindallia*, *Ercella* and *Anoxynatronum* were the primary bacterial genera and *Methanothrix*, *Methanobacterium*, *Nitrososphaera* and *Methanosarcina* were the dominant archaeal genera. Upon combining the microbial diversity and the geochemical properties of the two sample types, the microbial community containing *Desulfomicrobium*, *Paracoccus*, *Tindallia*, *Methanobacterium*, *Methanothrix* and *Nitrososphaera* was better adapted to the biodegradation of flotation collectors, and the microbial community consisting of *Paracoccus*, *Exiguobacterium*, *Methanothrix* and *Methanobacterium* was more efficient at hydrolyzed polyacrylamide (HPAM) biodegradation. In addition, a large proportion of unclassified OTUs has indicated that recycling effluent is a worthy resource for isolating new strains from the Firmicutes phylum.

## 1 Introduction

China has been the world’s leading producer of metallurgical-grade alumina over the last 16 years(1). The operational yield of the Chinese alumina industry in 2017 exceeded 69 million tons, with a year-on-year growth of 13.32%, which resulted in the rapid depletion of high A/S (mass ratio of alumina to silica) bauxite as well as a great challenge to Chinese bauxite resources(2). Bauxite desilication by flotation, which could be used for low A/S bauxite that is found in large reserves in China, became an important technical method for the sustainable development of the aluminum industry in China(3, 4)

A large amount of water is needed for the bauxite desilication process, and a large quantity of water can be saved by water recycling through flocculation and the rapid settlement of the pulp(5). However, the production index becomes worse with the increased circulation time of the circulating water; the periodic replacement of circulating water was the primary method for solving this problem in bauxite flotation plants, resulting in a problem of recycling effluents. These recycling effluents contained high concentrations of ions and the residues of beneficiation reagents, such as collectors and flocculants(6, 7). The periodic discharge of recycling effluents would have potential environmental risks, but little study has been devoted to this approach. However, with the rapid economic growth and social development in China, the associated problems of water contamination and shortages of water resources have become serious bottlenecks that challenge and limit sustainable development at the regional as well as national levels(8, 9). Finding an efficient and safe method for treating the recycling effluents, such as biological methods, is an urgent task.

Because of the importance of bauxite desilication by flotation for the alumina industry of China and with the increased use of water resources, the microbial communities in the recycling effluents from desilication processing were revealed and linked with the chemical and geographical conditions to create effective advice for the prevention and treatment of these problems. The compositions, diversity and dynamics of the given microorganisms may interfere in the efficiency and stability of wastewater treatment plants(10). Therefore, studies about the microbial community in these systems have been focused on the functions and performance of wastewater treatment(11). In this study, recycling effluents from the desilication process were collected before being discharged at the bauxite flotation plant in Henan province, and the community structures were assayed using 16S rRNA genes by high-throughput sequencing method.

## 2 Materials and methods

### 2.1 Site description and sample collection

Samples of recycling effluent were collected from the bauxite flotation plant located in Henan province. Recycling effluents of concentrates and tailings were collected from a circulating cistern (Fig 1), and they were named HNC and HNT, respectively.

**Fig 1.**
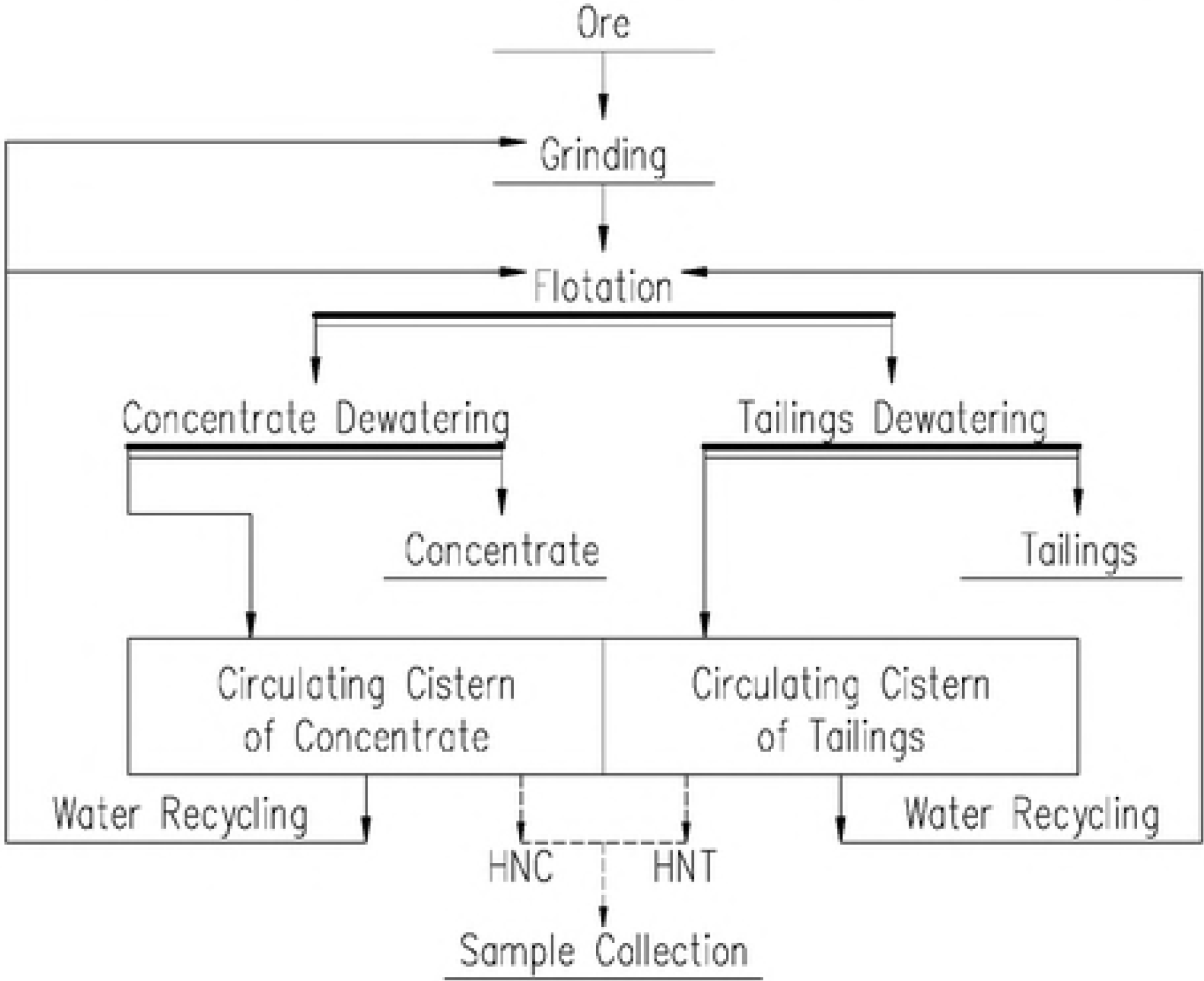
Flow chart of desilication process and sample collection

Water samples were collected in July of 2018. Approximately 25 L of recycling effluent from each circulating cistern was collected, and the temperature and pH of each sample were detected at the same time. The samples were then filtered through a 0.22 μm hyperfiltration membrane with a vacuum pump. The sediments on the membranes were then immediately transferred to a tube and stored at −20° for further molecular analysis. The filtered water samples were sent for chemical analysis.

### 2.2 Geochemical analysis

We measured the concentrations of 28 elements, including Al, As, Ca, Cr, Cu, Fe, K, Mn, Na, Pd, Zn and other elements, by Inductively Coupled Plasma Mass Spectroscopy (ICP-MS, G8403A, Japan).

### 2.3 DNA extraction, PCR and sequencing

The DNA in the water samples was extracted from 200 mg of sediment with an E.Z.N.A. Soil DNA Kit (OMEGA, USA). The DNA integrity was detected by agarose gel electrophoresis. Accurate quantitative control of the genomic DNA added to the PCR reaction system was achieved after the determination of the genomic DNA with a Qubit 3.0 DNA detection kit.

The V3-V4 region(12)of the 16S rRNA genes was amplified with the primer pair 341F (5′ -CCCTACACGACGCTCTTCCGATCTG (barcode) CCTACGGGNGGC WGCAGA-3′) and 806R (5′ -GACTGGAGTTCCTTGGCACCCGAGAATTCCAGACTACHVGGGTATCTAATCC-3′) combined with the universal primers on the MiSeq sequencing platform.

The PCR reactions of the first round had a total volume of 30 μl, which contained 10-20 ng of genomic DNA, 15 μl of 2×Taq master Mix, 1 μl of Bar-PCR primer F (10 μM), 1 μl of primer R (10 μM), and the remaining volume of H_2_O. The cycling conditions of the first round were 94° for 3 min, 5 cycles of 94° for 30 s, 45° for 20 s, and 65° for 30 s, 20 cycles of 94° for 20 s, 55° for 20 s, and 72° for 30 s, and a final extension at 72° for 5 min. The PCR reactions of the second round had a total volume of 30 μl, and they contained 20 ng of genomic DNA, 15 μl of 2×Taq master Mix, 1 μl of primer F (10 μM), 1 μl of primer R (10 μM), and the remaining volume of H_2_O. The cycling conditions of the second round were 95° for 3 min, 5 cycles of 94° for 20 s, 55° for 20 s, and 72° for 30 s, and a final extension at 72° for 5 min.

The PCR products were purified by Agencourt AMPure XP (Beckman, USA) and quantified accurately by Qubit 3.0 DNA detection kit. The PCR products of each sample were mixed to a final concentration of 20 pmol and then sequenced with a MiSeq benchtop sequencer (Illumina, San Diego, USA) for 2×300 bp paired-ends sequencing.

### 2.4 Sequence data processing and statistical analysis

The sequenced reads with perfect matches to barcodes were split into sample libraries, and the chimeras were high-quality trimmed using Usearch. The OTUs were classified at a 97% similarity level and rarefaction analyses were conducted using Mothur(13). The curves from the rarefaction analysis and rank abundance analysis were marked using the originally detected OTUs from R Project 3.2.

The sequences were first filtered through a BLAST search against the RDP database(14), with a common confidence threshold of 80–90%, and the satisfactory sequences were phylogenetically assigned to taxonomic classifications via an RDP naïve Bayesian classifier and Bergey’s taxonomy at a 0.03 distance(15). A dendrogram was constructed with data derived from a hierarchical cluster analysis of the beta diversity distance matrix and an unweighted pair group method with arithmetic mean (UPGMA) cluster analysis(16). Bray-Curtis ordination was used to calculate the distance between samples. A phylogenetic tree was constructed with Python software and functional gene compositions were calculated with PICRUSt 1.0.0 software(17). All the other analyses were performed in R 3.2 with the vegan packages v. 2.0-10.

## 3 Results and discussion

### 3.1 Geochemical properties of samples

The plant from which the samples were collected is located in Guanyinsi village, Henan province, China. The two samples were collected outdoors, from the exposed cisterns, and the colors of the two samples were light yellow with some suspended substances. The temperature of them both was approximately 31°C, and the pH values were 9.37 and 9.28 for the HNC and HNT samples, respectively. The detectable ion concentrations are listed in Table 1, and they showed similar concentrations in the two samples. The samples contained a very high level of Na from the addition of NaCO_3_ during bauxite flotation processing. The concentrations of Al, K and Ca were at a relatively higher level. In addition, the concentrations of Mg, Fe, Zn, Mo and Ba were at a relatively lower level. Some ions, such as Ti, As, Pb, Sb, B, Mn, and Se, showed very low concentrations, while some ions, such as Cu, Ni, Cd, Ag and others, were not detected. The organics in the samples were the residuals of collectors (such as hydroximic acid, oleic oil and others) and flocculants (primarily HPAM).

**Table 1.**
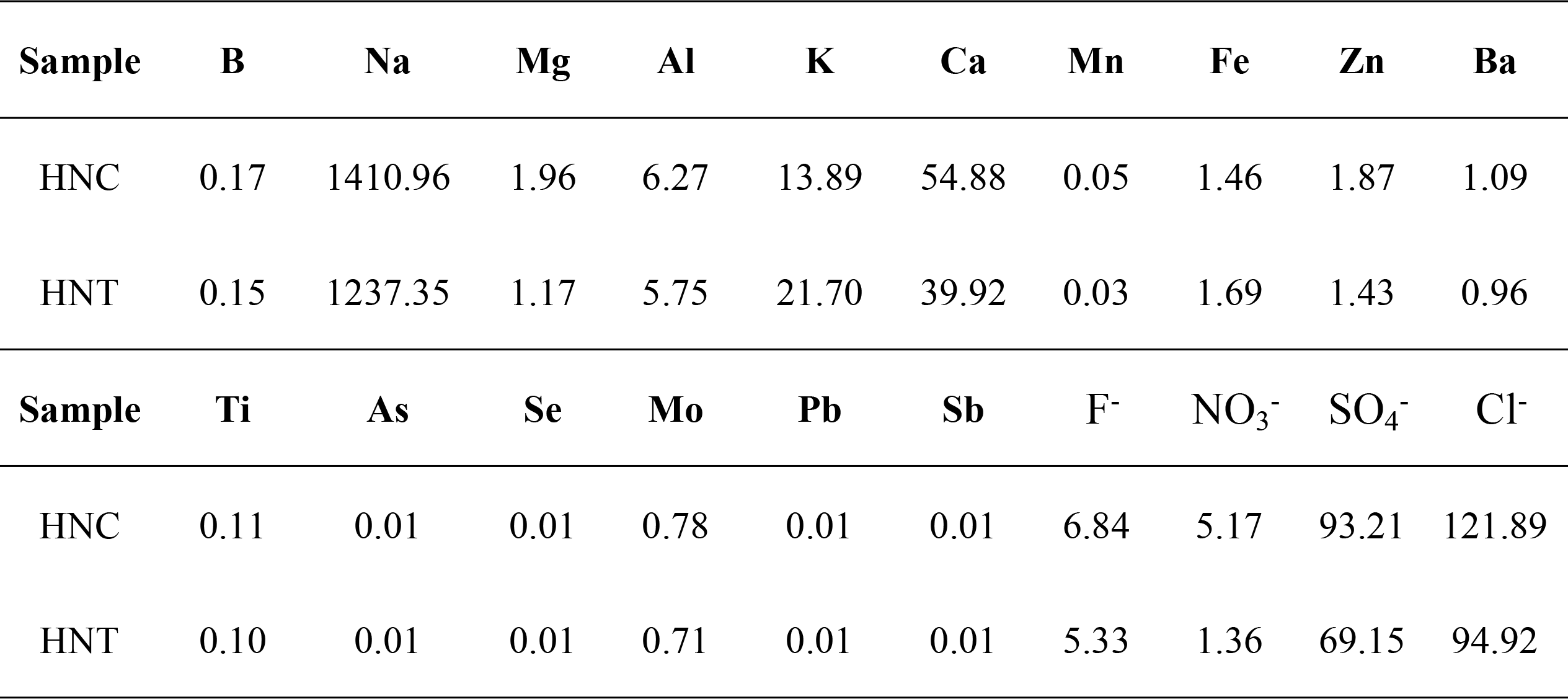
Geochemical properties of the water samples (mg/L)

| Sample | B | Na | Mg | Al | K | Ca | Mn | Fe | Zn | Ba |
| --- | --- | --- | --- | --- | --- | --- | --- | --- | --- | --- |
| HNC | 0.17 | 1410.96 | 1.96 | 6.27 | 13.89 | 54.88 | 0.05 | 1.46 | 1.87 | 1.09 |
| HNT | 0.15 | 1237.35 | 1.17 | 5.75 | 21.70 | 39.92 | 0.03 | 1.69 | 1.43 | 0.96 |
| Sample | Ti | As | Se | Mo | Pb | Sb | F<sup>-</sup> | NO<sub>3</sub><sup>-</sup> | SO<sub>4</sub><sup>-</sup> | Cl<sup>-</sup> |
| HNC | 0.11 | 0.01 | 0.01 | 0.78 | 0.01 | 0.01 | 6.84 | 5.17 | 93.21 | 121.89 |
| HNT | 0.10 | 0.01 | 0.01 | 0.71 | 0.01 | 0.01 | 5.33 | 1.36 | 69.15 | 94.92 |

In all, the two samples were both taken from the bauxite flotation process. They had similar ion concentrations, pH values, temperatures and flocculant concentrations, but they had different concentrations of collectors for the chemical adsorption on concentrate and no effect on the tailings.

### 3.2 Analyses of OTUs

Both sequencing data sets were separated from one another to investigate the core microbiome in recycling effluents of the concentrate and tailings. Furthermore, a differentiation between bacteria and archaea was made to avoid the loss of the archaeal OTUs due to their low sequence reads. More than 35000 high-quality sequences with at least 1600 OTUs for bacteria as well as more than 27000 high-quality sequences with at least 320 OTUs for archaea were used for the microbial community composition analysis of the two samples after filtering out the chimeras. The asymptotic level of the Shannon rarefaction curves, at a 97% similarity level, suggested that the number of OTUs were sufficient for this study (Fig 2). The Shannon diversity index of bacterial communities was significantly higher than that of the archaeal communities, which suggested that the bacterial communities were more varied.

**Fig 2.**
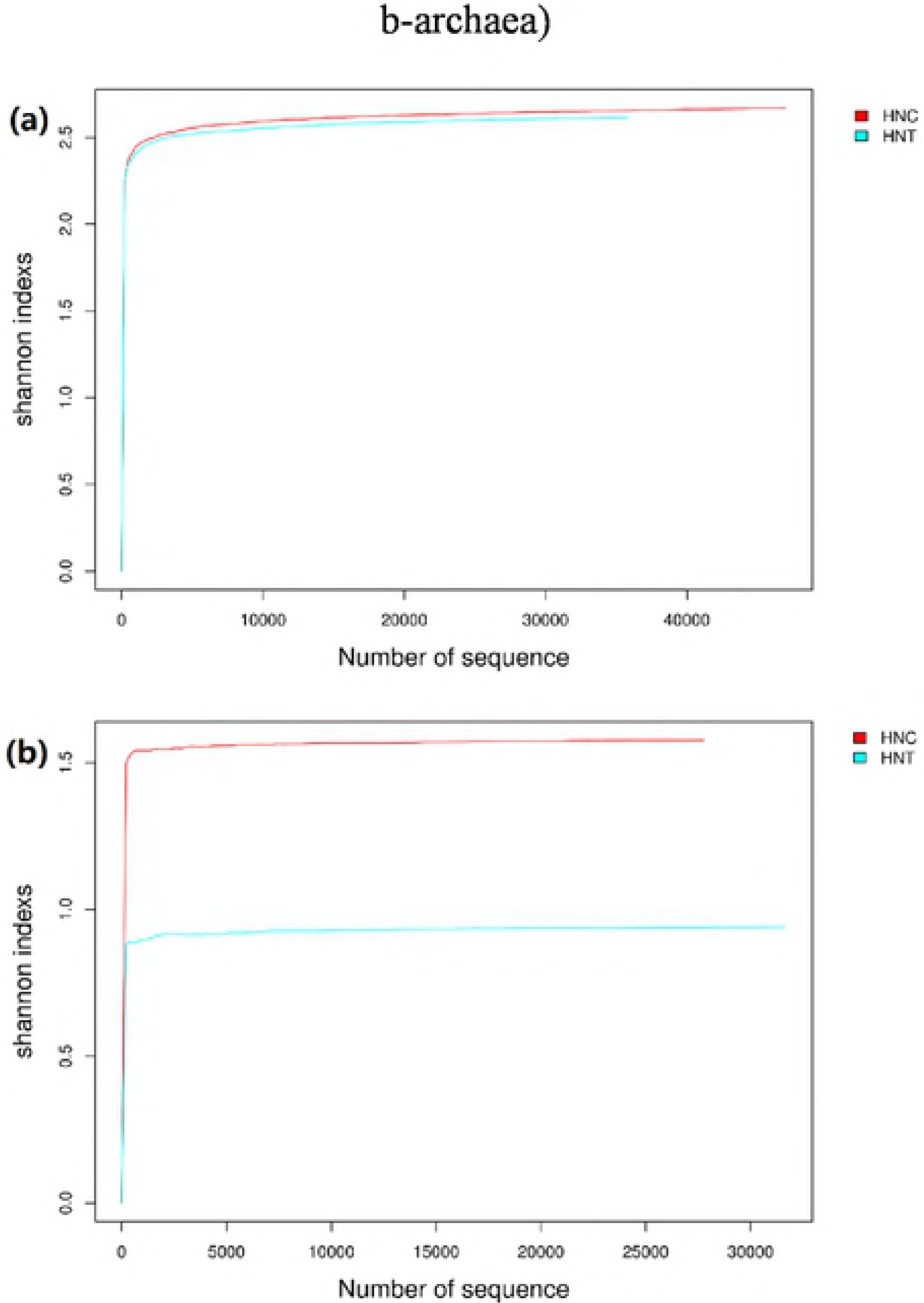
Curves from the Shannon rarefaction analysis (a-bacteria, b-archaea)

Rank-abundance curves are used to interpret two aspects of sample diversity, namely the richness and evenness of the species in the sample. Fig 3 shows that the samples had low diversity index for the narrow change along the lateral axis while the two sites had a low degree of uniformity for both archaeal and bacterial compositions because of the steep shapes of the curves. The rarefaction curves combined with the rank abundance curves suggested that the work was able to reveal the real microbial community compositions of the samples.

**Fig 3.**
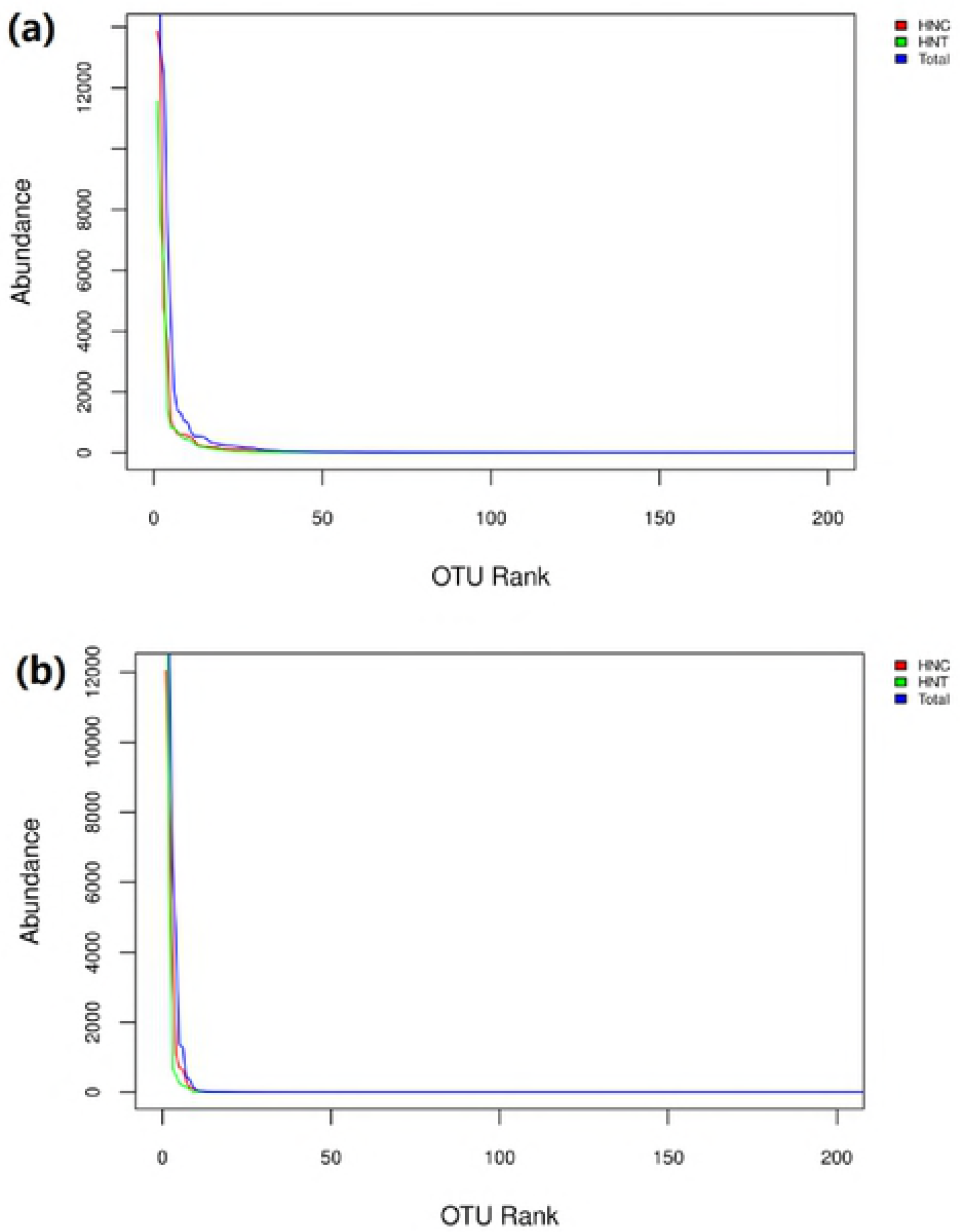
Rank abundance curves based on OTU ranks (a-bacteria, b-archaea)

A Venn diagram could reflect the similarity and overlap of the OTU composition between samples visually (Fig 4). The samples in this study had low similarity rates in both archaeal and bacteria communities at the OTU level. The bacterial OTUs were more abundant than archaeal OTUs in the two samples, and the microbial composition of the HNC were more abundant than that of the HNT. For both the archaeal and bacterial communities, the common OTUs both detected in the two samples were at a low level for each total OTU number, at 4.09% and 2.57%, respectively. That finding revealed the significant diversity between the HNC and HNT samples that might be caused by the different properties of concentrate and tailings, although both of them were involved in the water cycle of the bauxite flotation process.

**Fig 4.**
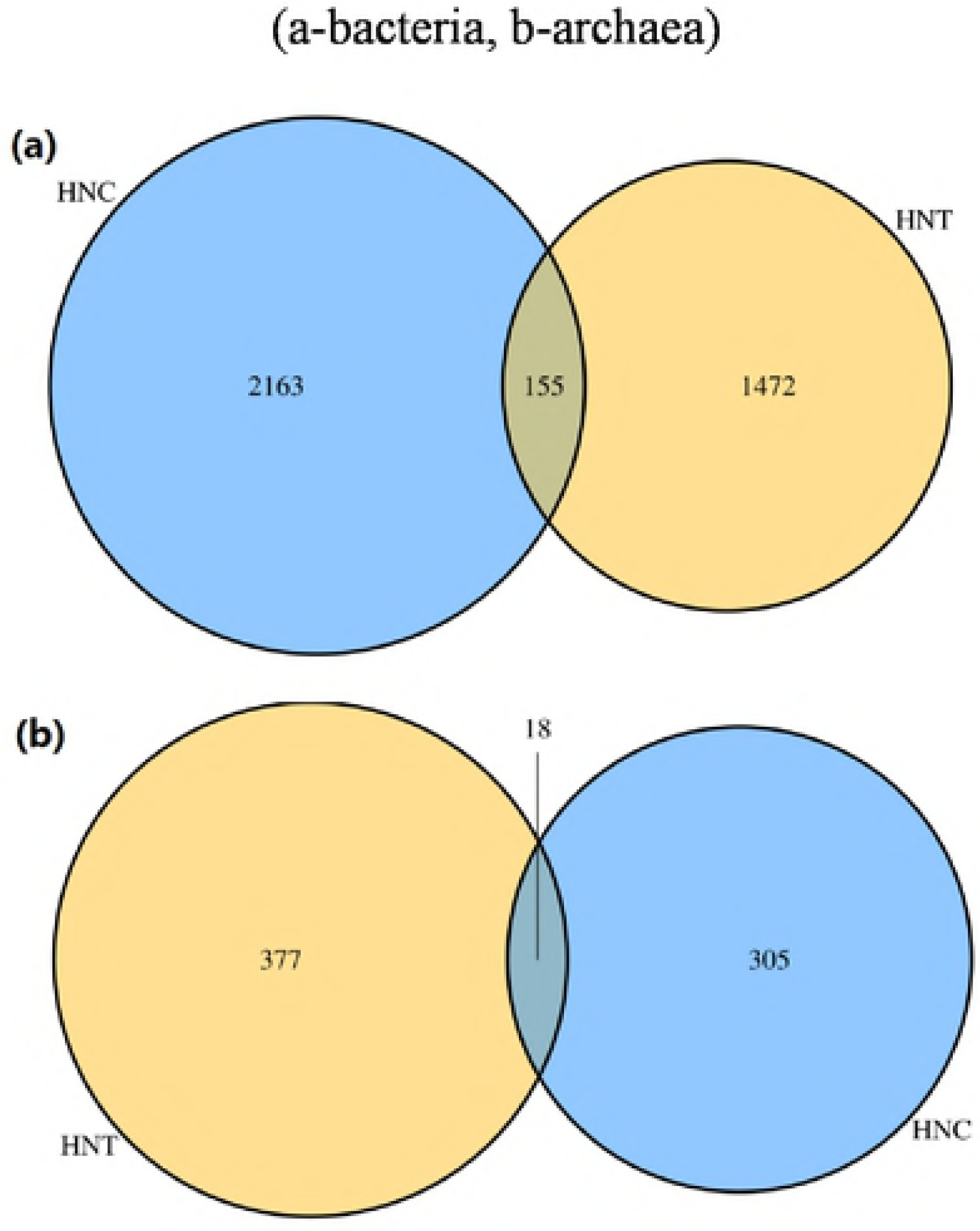
Similarity and overlap analysis of OTUs by Venn diagram (a-bacteria, b-archaea)

### 3.3 Analyses of microbial communities

To identify effective biological methods for the prevention and treatment of the recycling effluents problem, it is vital to comprehensively understand the involved microbial communities in terms of taxonomic compositions, similarity and diversity. A taxonomic composition analysis is one of the most frequently used bioinformatics analyses for microbial communities(18).

#### (1) Bacterial communities

As illustrated in Fig 5, the dominant bacterial genus was unclassified in the two samples accounting for 38.47% and 45.75% for HNC and HNT, respectively. Interestingly, almost all of the unclassified OTUs fell into the Firmicutes phylum, which suggested a new source for seeking new species in the Firmicutes phylum. For the HNC sample, with the exception of the unclassified bacterial genus, *Desulfomicrobium, Paracoccus* and *Tindallia* were the top three predominant genera, accounting for 29.62%, 10.17% and 8.01%, respectively, followed by *Exiguobacterium, Dethiosulfatibacter, Nitrincola, Anoxynatronum* and *Ercella* with a proportion over 1%. For the HNT sample, except for the unclassified bacterial genus, *Paracoccus* and *Exiguobacterium* were the top two predominant genera, accounting for 21.34% and 17.99%, respectively, followed by *Ercella, Tindallia, Pirellula, Anoxynatronum* and *Ercella*, with a proportion between 1% to 3%.

**Fig 5.**
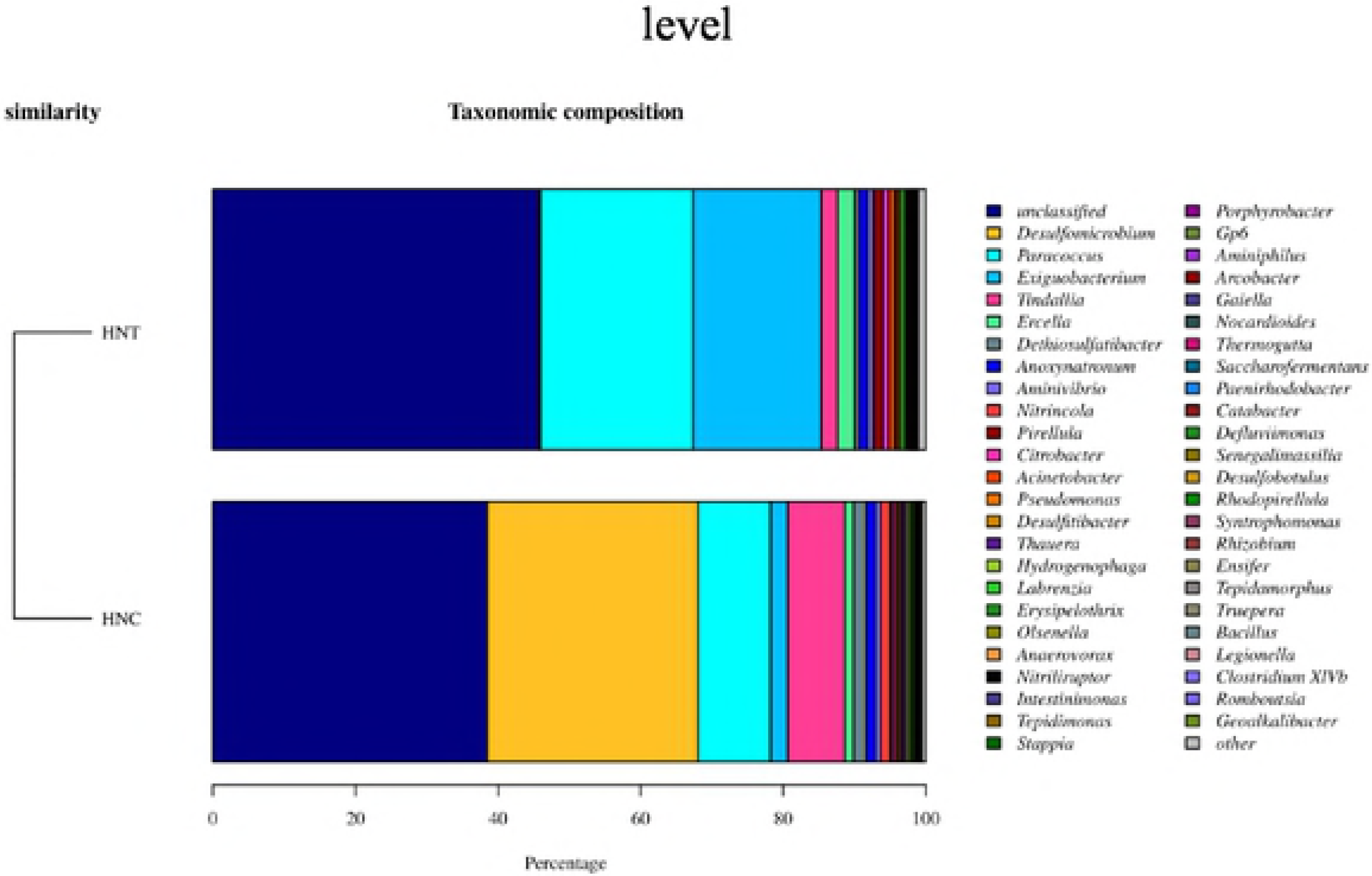
Taxonomic distribution of bacterial OTU abundance at the phylum level

According to the taxonomic comparison results, the dominant species were selected and the evolutionary relationships and abundance differences in the dominant microorganisms in the sequenced environmental samples were understood from the phylogenetic system (Fig 6). According to the phylogenetic analysis, 99.03% of the bacterial OTUs fell into the following six phylogenetic divisions: Proteobacteria, Firmicutes, Synergistetes, Planctomycetes, Actinobacteria and Acidobacteria. Proteobacteria and Firmicutes were the dominant phyla while the others contributed little to the bacterial community structures.

**Fig 6.**
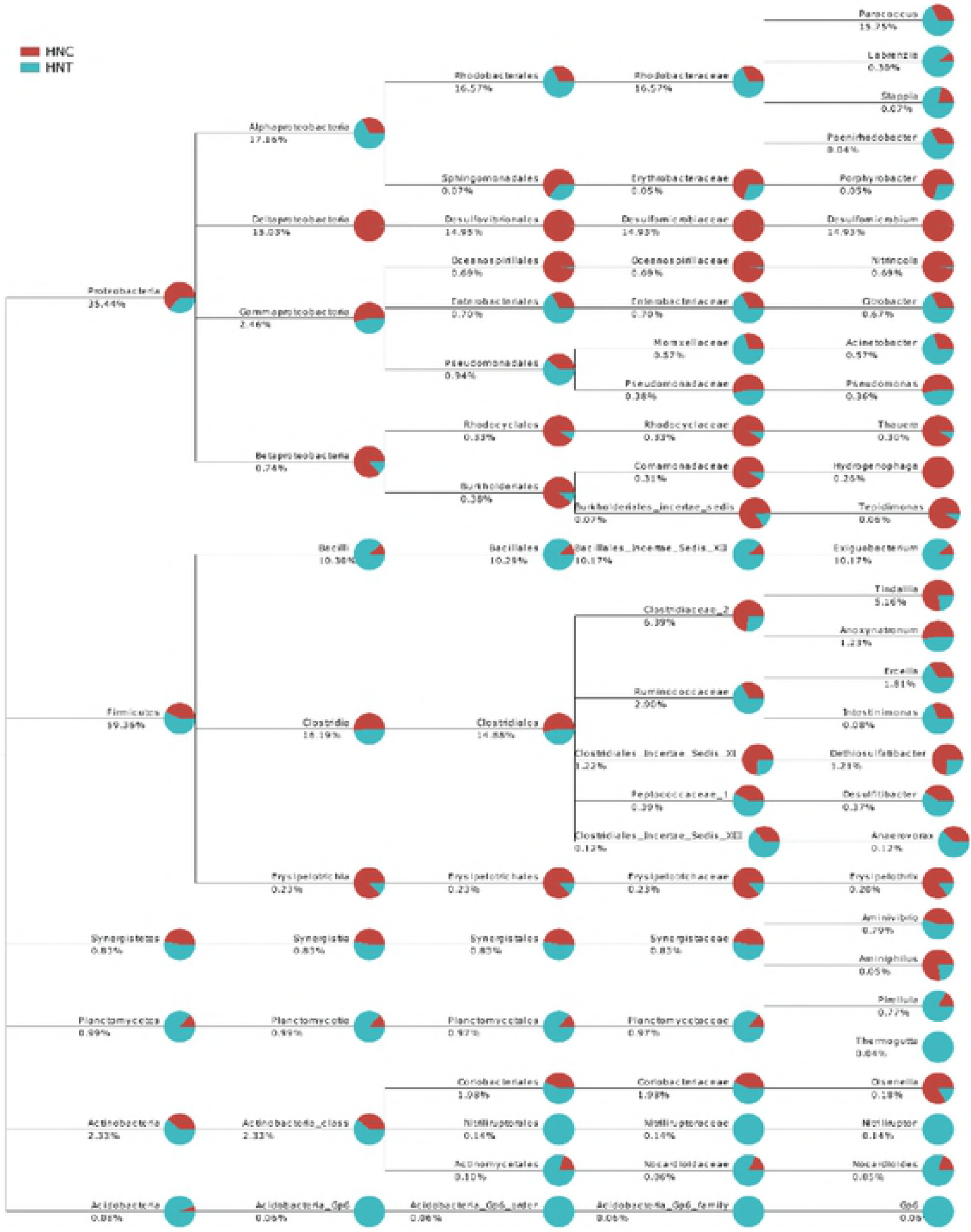
Phylogenetic compositions and community structures of bacteria

Within the Proteobacteria phylum, which accounted for 35.44% of the bacterial communities, *Paracoccus* belonged to Alphaproteobacteria (mostly found in HNC) and *Dethiosulfatibacter* belonged to Desulfomicrobium (mostly found in HNT) were the major genera. Firmicutes accounted for 59.36% of all the OTUs, but more than half of that OTUs were classified into unknown genera, and 10.17% of all the OTUs fell into the *Exiguobacterium* genus while 5.16% fell into *Tindallia*.

*Paracoccus* was found in both samples and played an important role in the bacterial community structures. HPAM, hydroximic acid and oleic oil were the primary organic pollutants in recycling effluents, and the *Paracoccus* genus emerged as a versatile species for the bioremediation of various recalcitrant pollutants, such as pyridine(19), dimethylformamide(20), N-methylpyrrolidone(21), and others. *Paracoccus* was also reported to not only reduce nitrate under an aerobic atmosphere but also to convert ammonium to nitrogen gas via hydroxylamine, nitrite, nitrate and nitrous oxide, which indicated that another nitrogen removal pathway immediately accessible via hydroxylamine without any accumulation of nitrate and nitrite could be responsible for this phenomenon(22). Hydroxylamine from organic pollutants and nitrate from recycling effluents were the primary nitrogen source for microbial growth as well as biodegradation actions.

Although several OTUs were detected in the HNT sample, *Desulfomicrobium* was in the majority in the HNC. *Desulfomicrobium* is gram-negative and is commonly found in aquatic environments with high levels of organic matter as well as in sewage sludge as a major community member of extreme oligotrophic habitats(11). Reports revealed the good bioremediation potential of the reductive process performed by *Desulfomicrobium*, and the degradation pathway was that sulfate and other oxidized sulfur compounds serve as terminal electron acceptors and are reduced to H_2_S(23). The two samples in this study had similar sulfate concentrations as well as temperature and pH, and the diversity in *Desulfomicrobium* between two samples might be due to that residue of hydroximic acid and oleic oil mostly in concentrate pulp while little of that was associated with tailing pulp for collectors, which were compatible with concentrate(24).

Contrary to the distribution of *Desulfomicrobium*, there was little *Exiguobacterium* in the HNC but quite a lot in the HNT. The genus *Exiguobacterium*, which belongs to the Bacilli class, is characterized by the following features: the cells are gram-positive, non-endospore-forming, facultatively anaerobic and oxidase-negative (25). *Exiguobacterium* has been reported to promote plant growth and the degradation of environmental pollutants, to be halotolerant(26, 27) and to be able to produce highly effective proteolytic enzymes(28). The HNT sample had a pH value of approximately 9.28, which provided an environment for the bacteria belonging to the *Exiguobacterium* genus to show their applicability to the biological neutralization of alkaline wastewater(25). The HNC also had a pH value of 9.37 but little *Exiguobacterium*, perhaps because of the inhibition of carbon sources (such as hydroximic acid and oleic oil) and the competition from *Desulfomicrobium*.

*Tindallia, Anoxynatronum* and *Ercella* were all detected in the two samples as auxiliary members, and they were all reported to be alkaliphilic and good for potential use in treating alkaline wastewater(29–31).

#### (2) Archaeal communities

Archaea, which were originally found in extreme environments, were known to be ubiquitous and crucial partners in numerous microbiomes, either in environmental settings or associated with holobionts(32), and they were no exception in this research. Fig 7 reveals that *Methanobacterium* and *Methanothrix* were the dominant archaeal genera in the two samples, accounting for 46.67% and 30.24%, 75.13% and 19.99% for HNC and HNT, respectively. *Nitrososphaera* was also abundant in HNC, with a proportion of 18.62%, while it contributed only 4.17% to HNT. *Methanosarcina* was found in both samples and accounted for a low proportion of the total.

**Fig 7.**
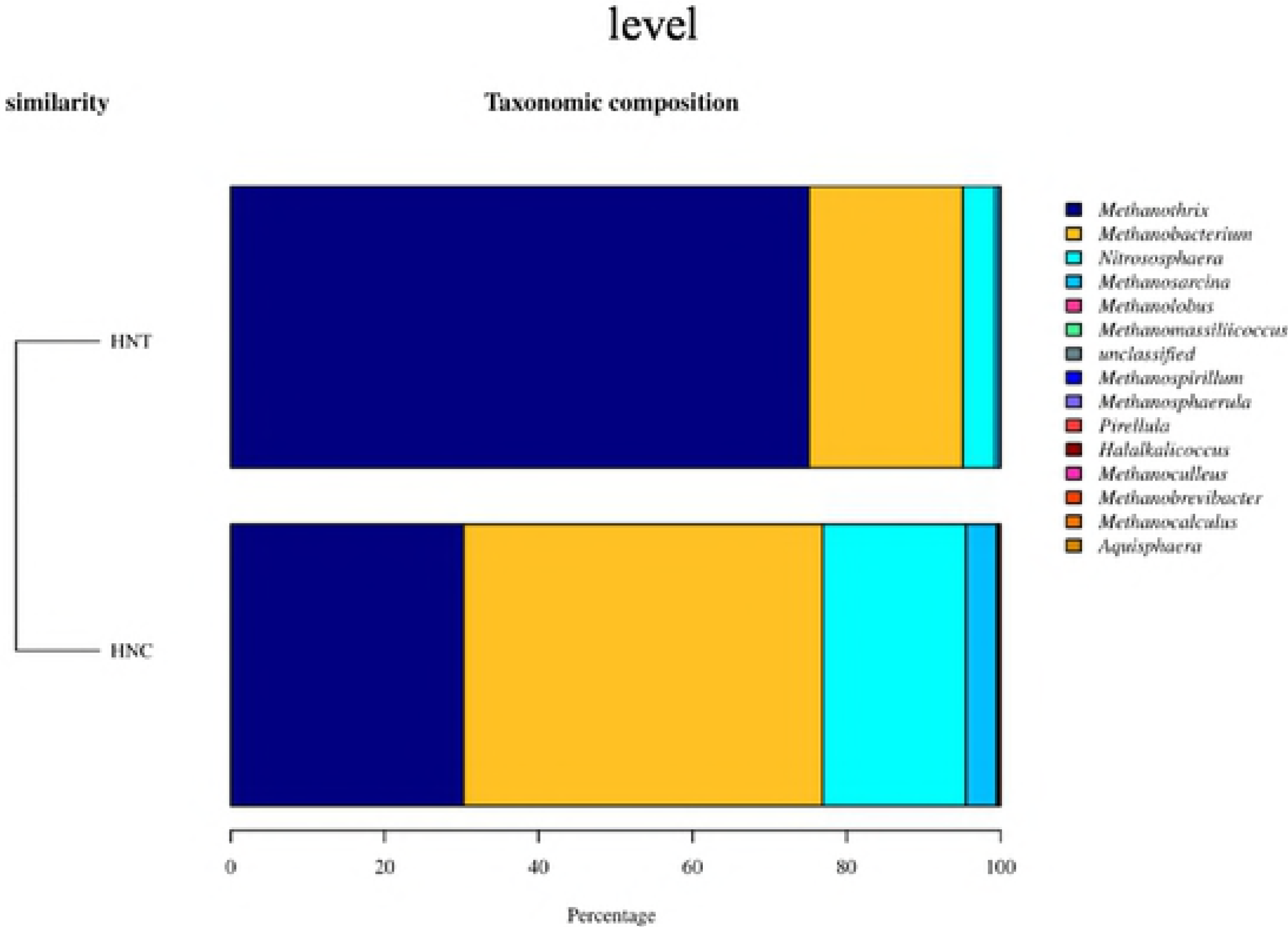
Taxonomic distribution of archaeal OTU abundance at the phylum level

According to the phylogenetic analyses (Fig 8), almost all of the archaeal OTUs fell into the three archaeal phyla Euryarchaeota, Thaumarchaeota and Planctomycetes. Euryarchaeota was dominant and Planctomycetes was seldom identified. Euryarchaeota contains methane-forming, extremely halophilic, sulfate-reducing, and extremely thermophilic sulfur-metabolizing species(33). Although there was a similar distribution in the two samples at the phylum level, *Methanobacterium* was more abundant in the HNC sample and *Methanothrix* was ascendant in the HNT at the genus level under the Euryarchaeota phylum.

**Fig 8.**
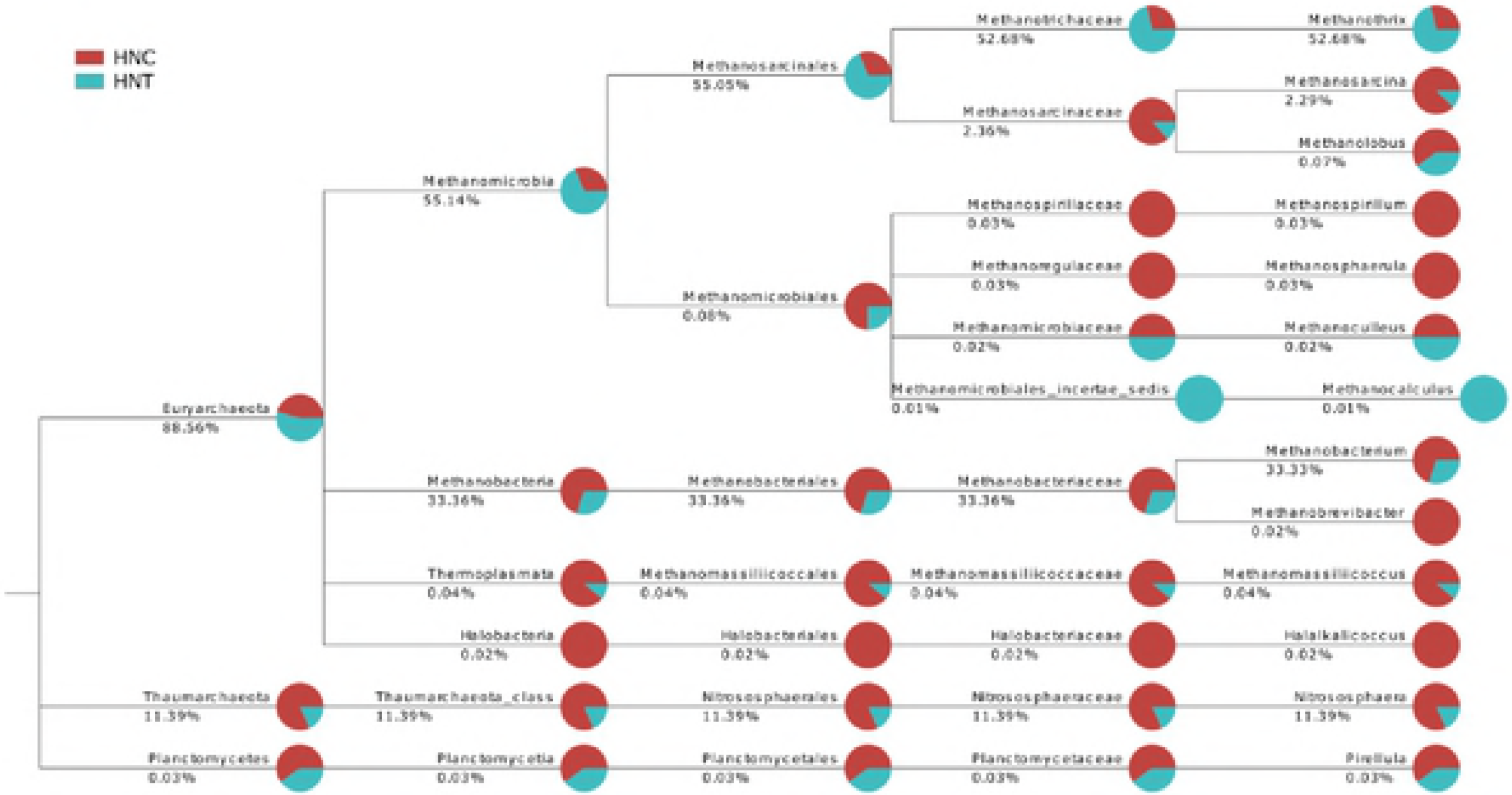
Phylogenetic compositions and community structures of archaea

*Methanothrix*, a genus of Methanosarcinales that are typical acetoclastic methanogenic archaea(34), made up more than half of all the OTUs, while *Methanobacterium*, a genus of Methanobacteriales that could reduce CO using molecular hydrogen (hydrogenotrophic) and produce methane(35), accounted for approximately one third of all the OTUs. These two primary genera belong to the methanogenic archaea and made up more than 85% of the archaeal communities. The methanogens were the initially and most widely recognized archaea, and they were considered to play a prominent role in the degradation of complex organic compounds by a consortia of anaerobic microorganisms. The methanogens are dependent on sodium ions for methane formation(36) and the sodium ion concentrations were at a high level (more than 1200 mg/L, as shown in Table 1) in the samples, which provided an appropriate environment for the high abundance of methanogenic archaea.

It is unclear how archaea communicated or structurally interacted with their environment. Thus, the methanogens certainly represented the keystone species, but the archaeal diversity beyond the methanogens must not be neglected. *Nitrososphaera* and *Methanosarcina* are the remaining genera found in these samples, and unclassified genera made up less than 0.1%, which verified the low diversity of archaea again. *Nitrososphaera*, which is one of the primary phylotypes of ammonia-oxidizing archaea (AOA)(37), had also been directly linked to active nitrification; it was found that the biotransformation of micropollutants were involved in the co=metabolism of ammonia-oxidizing microorganisms(38, 39). *Methanosarcina* is one of the few methanogens shown to participate in direct interspecies electron transfer (DIET) by directly receiving electrons to reduce CO_2_ into CH_4_(40), and it appeared to be of crucial importance to the anaerobic digestion process(41).

#### (3) Microbial community diversity between two samples

Although the syntrophic and competitive relationships between archaea and bacteria remained completely unclear, a consortia of bacterial and archaeal communities played an important role in the degradation of complex organic compounds(36). The ways in which archaea and bacteria define their niche within a complex setting could be revealed by extensive and in-depth studies on microbial community diversities in different environments.

The similarity of the pH values, water temperature, ion concentrations and types of organic pollutants between HNC and HNT gave an intimation of the similarity in components in both the archaeal and bacterial communities. The diversity in the concentrations of collectors was the primary reason for the diversity of distributions in the microbial composition. The primary bacterial compositions of both samples consisted of *Desulfomicrobium, Paracoccus, Tindallia, Methanobacterium, Methanothrix* and *Nitrososphaera*, which were all reported as alkaliphilic bacteria for biological neutralization(29, 30, 42, 43). The primary archaeal compositions in both samples consisted of (the genera *Methanothrix, Methanobacterium* and *Methanosarcina*) and AOA (genus *Nitrososphaera*), which appear to be part of a key strategy to adapt to a complex environment(32).

According to the sample HNC result, the community primarily made up of *Desulfomicrobium*, *Paracoccus*, *Tindallia*, *Methanothrix*, *Methanobacterium* and *Nitrososphaera* seemed better adapted to the biodegradation of the flotation collectors. Flotation collectors are small molecular organic materials, and they could be degraded by bacteria more easily. The sulfate reduction and ammonia oxidization tasks were performed by *Desulfomicrobium* and *Nitrososphaera*, while other genera played a supporting role.

According to the result for the HNT, the community primarily made up of *Paracoccus, Exiguobacterium, Methanothrix* and *Methanobacterium* was considered to be more efficient at biodegrading the flocculant. HPAM, the flocculant in the samples, was a bio-refractory polymer compound with amide groups and a carbon backbone that could be the energy source for microorganisms. In these habitats and with flocculant as a substrate, *Exiguobacterium* produced lactate, acetate and formate, and these products were converted into methane by *Methanothrix* and *Methanobacterium*. *Paracoccus*, the dominant bacterial genus, could grow under aerobic and denitrifying conditions, and it showed great potential at hydrogen sulfide removal from synthetic biogas(44, 45). *Exiguobacterium* was reported to grow through the utilization of molecules containing acyl groups(46). These two bacterial genera supported methanogenesis by methanogens, which was important at the end of the anaerobic digestion chain for biomass conversion.

## 4 Conclusions

The archaeal and bacterial communities in recycling effluents from bauxite flotation plant were revealed, and the diversity between the recycling effluent of concentrate and that of tailings was studied. Of the unclassified OTUs at the genus level, almost all fell into the Firmicutes phylum, and they accounted for approximately 40 percent of all the OTUs. Therefore, this recycling effluent was a worthy resource for the isolation of new strains in the Firmicutes phylum. At the genus level, *Paracoccus*, *Desulfomicrobium*, *Exiguobacterium*, *Tindallia*, *Ercella* and *Anoxynatronum* were the primary members of the bacterial community, while *Methanothrix, Methanobacterium, Nitrososphaera* and *Methanosarcina* dominated the archaeal community. The biodegradation of the pollutants in the recycling effluents involved the synergism of bacteria and archaea. Combined with the geochemical properties of recycle effluents, the microbial community containing *Desulfomicrobium, Paracoccus, Tindallia, Methanobacterium, Methanothrix* and *Nitrososphaera* was more adept at the biodegradation of flotation collectors, such as hydroximic acid and oleic oil. The microbial community consisting of *Paracoccus*, *Exiguobacterium*, *Methanothrix* and *Methanobacterium* was more efficient at the biodegradation of HPAM. The results of this study provided a practical orientation on the biotreatment of recycling effluents from the bauxite flotation process.

## Acknowledge

This research was supported by Funds of the Research and Innovation Project of Graduate Students of Central South University (1053320170205), Fundamental Research Funds for the Central Universities of Central South University (502211704), State Key Laboratory of Advanced Technologies for Comprehensive Utilization of Platinum Metals (SKL-SPM-201809), State Key Laboratory of Applied Microbiology Southern China (SKLAM005-2016), National Natural Science Foundation of China (51320105006, 51504106 and 51871250), Science and Technology Project of Yunnan (2015FB204)

